# Hearing elliptic movements reveals the imprint of action on prototypical geometries

**DOI:** 10.1101/2022.10.30.514456

**Authors:** Etienne Thoret, Mitsuko Aramaki, Lionel Bringoux, Sølvi Ystad, Richard Kronland-Martinet

## Abstract

Within certain categories of geometric shapes, prototypical exemplars that best characterize the category have been evidenced. These geometric prototypes are classically identified through the visual and haptic perception or motor production and are usually characterized by their spatial dimension. However, whether prototypes can be recalled through the auditory channel has not been formally investigated. Here we address this question by using auditory cues issued from timbre-modulated friction sounds evoking human drawing elliptic movements. Since non-spatial auditory cues were previously found useful for discriminating distinct geometric shapes such as circles or ellipses, it is hypothesized that sound dynamics alone can evoke shapes such as an exemplary ellipse. Four experiments were conducted and altogether revealed that a common elliptic prototype emerges from auditory, visual, and motor modalities. This finding supports the hypothesis of a common coding of geometric shapes according to biological rules with a prominent role of sensory-motor contingencies in the emergence of such prototypical geometry.

## Introduction

Our perceptual system stores and categorizes objects from our surroundings around canonical items, called prototypes, that are the most representative of their category. Geometric prototypes have been viewed as mainly emerging from visual experiences (Rosch, 1973; Rosch & Mervis, 1975; Feldman, 2000; Kalénine et al. 2011; Theurel et al. 2012), via the haptic sensory channel (Theurel et al., 2012; Kalénine et al. 2011) and via a motor restitution (Feldman, 2000; Kalénine et al., 2013). Alternatively, the possibility to recall prototypes through visual and haptic modalities or motor output may suggest common processing of these specific shapes. Whether such prototypes only rely on spatial cues that are present in the visual, haptic, and kinetic domains is still questioned. Could such geometric prototypes also be recalled through auditory stimuli that are solely based on dynamic motor cues? Answering this question would provide a new path to understanding the influence of the motor system on human learning, memory, and cognition. This would indeed suggest that human movement dynamics and more generally the sensory-motor contingencies influence the structuration of geometric shape categories during development.

Interestingly, studies revealed the proficient role of the auditory modality to perceive movements and shapes through timbre variations of monophonic sounds (Merer et al. 2008; 2013). More strikingly, Thoret et al. (2014) demonstrated that auditors listening to synthesized monophonic friction sounds corresponding to those produced by the pencil of someone drawing on a paper, were able to recognize specific kinematics characterizing biological motion, in particular the 1/3 power law linking the tangential velocity of the hand movement to the curvature of the drawn shape (Lacquaniti et al., 1983). In addition, they were even able to discriminate geometric shapes, such as a circle, an ellipse, and a line, simply by listening to synthetic friction sounds in which timbre variations revealed the velocity profile of the drawing movement. In follow-up studies, we demonstrated that these acoustic variations may even distort the visuomotor coupling of biological motions (Thoret et al, 2016a, 2016b). Taken together, these studies suggest that simple geometric shapes can be evoked through the auditory channel and that the auditory modality may play a significant role in the perception and production of geometric shapes.

Here we investigated the hypothesis that geometrical prototypes can be recalled through the auditory modality employing timbre variations of friction sounds evoking the velocity of biological movements. We focused on a particular shape category, ellipses, that encompass any closed shape of a conic section contained between a line and a circle. From a geometric point of view, an ellipse is principally described by its eccentricity representing its flatness. The eccentricity is a number comprised between 0 and 1: the flatter the ellipse, the higher the eccentricity. The line and the circle are two specific cases which eccentricities equal 1 and 0 respectively. From a dynamic point of view and concerning the 1/3 power law, the accelerations of an elliptic movement increase as the distance between the focal points of the ellipse increases (i.e., when the ellipse tends towards a line). Ellipses can be distinguished by ear from circles and lines and do not involve discontinuity movements. Hence, this geometric shape has been chosen for the present study.

Based on the dynamic model of biological motion described below, four experiments were designed to examine how participants assessed the prototypical ellipse through different modalities. The first three experiments aimed at highlighting the geometric prototype from visual and motor restitutions. The fourth experiment was the cornerstone of this series of experiments and focused on sounds’ ability to evoke an elliptic prototype based on the kinematics underlying the drawn ellipse. Consistency between this auditory prototypical ellipse and the visual and motor prototypes would support a common encoding of prototypical shapes.

Before presenting the four experiments, the dynamic model of elliptic motions used in the four experiments will be described in the following section. The results of these experiments are presented together.

### A model of biological elliptic motor dynamics

Biomechanical mechanisms involved in graphical production as handwriting have been extensively investigated. It has been shown that dynamic and geometric properties of elliptic motions can be modeled by explicit equations. The dynamic approach of movement production supports the idea that planar hand movements can be modeled by two harmonic oscillators (x(t), y(t)) whose frequencies, amplitudes and phases evolve over time (Hollerbach, 1981) with the following system:

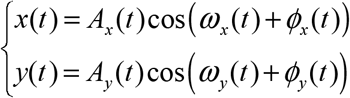

where A_x_ and A_y_ are the amplitudes, and ω_x_ and ω_y_ the frequencies, and ϕ_x_ and ϕ_y_ the phases of the two oscillators. In the case of periodic elliptic motions, this model can be simplified by equaling the amplitudes and frequencies of the oscillators (A(t) = A_x_ = A_y_ = A and ω(t) = ω_x_ = ω_y_ = ω):

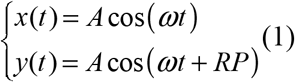

where A is the amplitude, 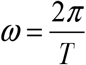 the frequency with T the period of the motion, and RP = ϕ_x_

- ϕ_y_ the relative phase between the oscillators *x*(*t*) and *y*(*t*).

Hence, this system co-defines the motion dynamics and the geometry, but may also be used to parameterize only the geometry of the entire ellipse whether or not the ellipse is considered dynamically. Practically, the eccentricity *e* of the ellipse and the relative phase RP are linked by the following relations demonstrated in Appendix A:

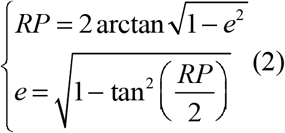

It should be noted that this model complies with the biological rules, namely the 1/3 power law (Lacquaniti et al., 1983), as demonstrated in Appendix A.

This model then enables the generation of biological elliptic motions whose trajectory can be continuously morphed from a line (RP = 0°) to a circle (RP = 90°) by simply acting on RP (see Figure 1). It will be used in the four following experiments in order to generate visual and auditory stimuli complying with biological motion. Motor productions of Experiment 2 will also be analyzed regarding this model.

**Figure 1.**
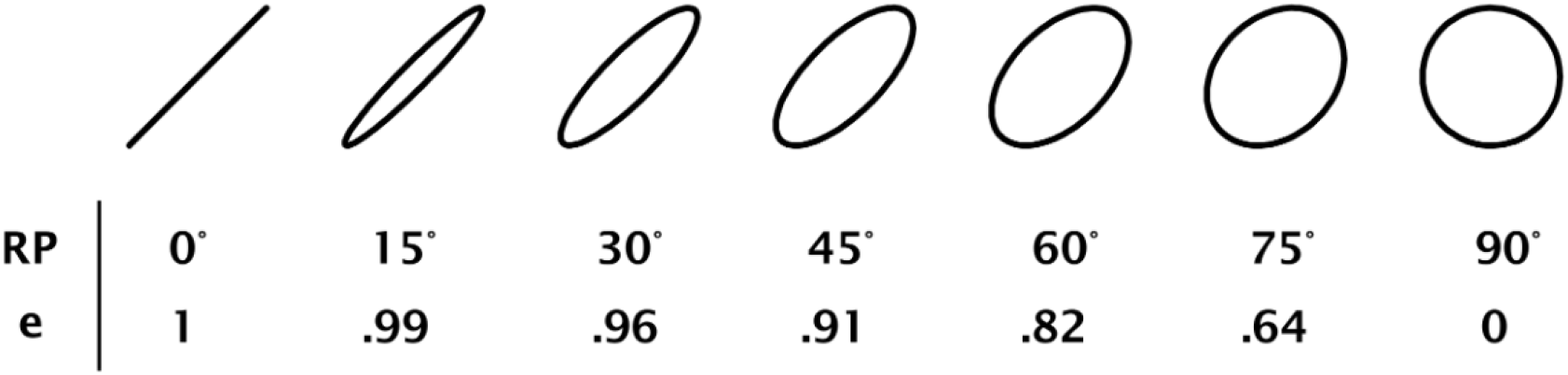
Continuum of different ellipse shapes from a line (left) to a circle (right) with the corresponding relative phases RP and eccentricities e.

Finally, the tangential velocity profile was generated according to the equations (1) and can be explicitly written as follows:

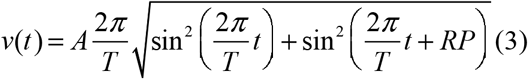

Although the mapping was arbitrarily fixed during the experiment, it is noticeable that it also varied according to the period T and the amplitude A of the motion.

### Experiment 1 – Static Visual Output

The first experiment aimed to evaluate the prototype of the ellipse from visual restitution (i.e., visual output). To that aim, an adjustment protocol was established. Participants were asked to adjust the eccentricity of static ellipses on a screen to evoke the most representative shape of this geometric category.

#### Methods

##### Participants

Twenty right-handed participants (8 women) with an average age of 30.6 years (SD = 12.8) voluntarily took part in the experiment. All the participants were naive to the purpose of the experiment. As in all the following experiments, participants gave their informed consent before the study, and the experiment was approved by the Aix-Marseille University ethical committee. For this experiment and the three subsequent ones, participants were selected from various groups, including personal acquaintances, students, engineers, researchers, and professors at the CNRS Campus Joseph Aiguier in Marseille, as well as individuals from the Sport Science Faculty of Aix-Marseille University in Marseille, France.

##### Stimuli

The visual stimuli were white static ellipses displayed on a black background with different sizes and eccentricities according to the dynamic model previously introduced (equations (1)). Three sizes defined by the amplitudes A_1_ = 3cm, A_2_ = 5cm, A_3_ = 10cm were chosen. The ellipses were rotated counterclockwise by 45° to conform to the preferential drawing inclination of an ellipse for right-handed persons (Danna et al., 2011).

##### Apparatus

The participants sat in front a computer screen (DELL 1907fp) with a resolution of 1280 × 1024 pixels and a frame rate of 60 Hz. The ellipses were displayed at the center of the screen and the interface was programmed with Max/MSP software (http://cycling74.com/). Participants modified the ellipse eccentricities by using a MIDI AKAY MPK keyboard. The experiment took place in a lighted room.

##### Task

The participants were asked to adjust the eccentricity of the static ellipse displayed on the screen so as to set the most representative geometric shape of this category. The notion of the most representative ellipse was explained in French to the participants by the following sentences (here translated in English): *Ellipses correspond to any closed shape between a line and a circle. When you imagine an ellipse, you may have one particular elliptic shape in mind. You will adjust the ellipse on the screen to display the ellipse you think is the most prototypical*. The adjustment protocol was based on the one proposed by Carlyon et al. (2010). The ellipse eccentricity was adjusted thanks to 6 different keys defined on the keyboard: “<<<” – “<<” – “<” and “>” – “>>” – “>>>”. Depending on the selected key, the ellipse eccentricity was modified with different step sizes. The higher the number of arrows the greater the step size, *i*.*e*. the modification of the eccentricity. To avoid a possible non-sensorial bias due to the orientation of the arrows (right or left), the action of the keys on the eccentricity differed across participants. Hence, for half of the participants, arrows that pointed to the right increased the eccentricity and for the other half they had the opposite effect. In addition, the step sizes were not the same in both directions. Hence, for a given participant, all the arrows in one direction (right or left) increased for instance the eccentricity by steps of {.01; .02; .03} while the others (left or right) decreased the eccentricity by steps of {-.1; -.001; -.005}. Finally, 11 repetitions for each size starting from 11 different initial eccentricities equally distributed between 0 and 1 were executed. The experiment was then composed of 33 trials, i.e. 3 {Sizes} x 11 {repetitions}, presented according to two pseudo-random series counterbalanced across the participants. The participants were prompted to explore the whole range of possibilities with the large arrows and to refine their adjustment with the smaller ones. For each participant, 33 final eccentricities were collected and the median was computed.

### Experiment 2 – Motor Output

The goal of this experiment was to evaluate whether a prototypical eccentricity of an ellipse can be elicited by a motor output. A motor production task was set up during which the participants were asked to draw the ellipse which best represents this geometric shape category.

#### Methods

##### Participants

Twenty participants (2 left-handed - 5 women) with an average age of 39.6 years (SD = 11.5) voluntary took part in the experiment and were naive to the purpose of the study. None of them participated in Experiment 1.

##### Apparatus

The participants sat in front of a Wacom Intuos5 graphic tablet enabling to record graphic movements with a spatial precision of 5.10^−3^mm and a sample rate of 129 Hz. The data were recorded and collected using an interface programmed with the Max/MSP software. The experiment was conducted in a lighted room.

##### Task

The participants were asked to repeatedly and continuously draw the most representative ellipse of this geometric shape category on the graphic tablet during 50 seconds. The notion of the most representative ellipse was explained with the same sentence as in Experiment 1. Participants saw their hands during the recording, but no trace was visible on the graphic tablet. The experiment comprised three sessions of 50 seconds. The participants were asked to draw *small, intermediate*, and *large* ellipses in two different orders counterbalanced across participants: 1) small – 2) intermediate – 3) large, or conversely. No template of the ellipses was presented, but the participants could train in advance by drawing the 3 different ellipses on the graphic tablet during a session preceding the experiment.

##### Data analysis

For each participant, recordings of the sampled coordinates (*x*(*t*), *y*(*t*)) of the stylus on the graphic tablet were collected. To eliminate the numerical noise, the raw data were smoothed with a Savitsky-Golay filter (Savitsky and Golay, 1964) with a 43 samples window and a 3^rd^-order interpolation. This is equivalent to a low-pass filter with a cutoff frequency of 8 Hz. A high-pass filter (Butterworth) with a cutoff frequency of .2 Hz was also applied to eliminate the spatial drift of the participants’ hands during their movements.

The geometric characteristics of the drawn ellipses were analyzed according to the relative phase. Computing the Hilbert transforms of *x*(*t*) and *y*(*t*) enabled to estimate the relative phase RP between two oscillators (Panter, 1965; Smith and Mersereau, 1991) with the following formula: 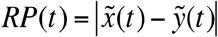, where 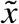 and 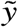 are the unwrapped Hilbert transforms of *x* and *y*. The median of the relative phases RP were then computed for each recording and then transformed into eccentricity. For each participant, 3 final eccentricities were collected and the median was computed.

### Experiment 3 – Dynamic Visual Output

Here, the Experiment 1 was reproduced with dynamic visual stimuli. The participants were asked to calibrate the eccentricity of the elliptic trajectory of a moving spotlight to create the most representative ellipse.

#### Methods

##### Participants

Twenty participants (1 left-handed; 3 women) with an average age of 32 years (SD = 11.4) voluntary took part in the experiment. None of these participants took part in Experiments 1 & 2.

##### Stimuli

The visual stimuli were white spotlights displayed on a black background moving on elliptic paths of different sizes and with different periods according to the dynamic model defined in equation (1). As in Experiment 1, the elliptic paths were rotated counterclockwise by 45°. Three periods of the spotlight motion (T_1_ = 1.2s, T_2_ = 1.5s, T_3_ = 1.8s) inducing different spotlight speeds were chosen.

##### Task

The adjustment protocol and the task were the same as in Experiment 1. For each (size) x (period) pair, 11 repetitions were performed starting from 11 different initial eccentricity values equally distributed between 0 and 1. The experiment was composed of 99 trials, i.e. 3 {sizes} x 3 {periods} x 11 {repetitions}, presented according to two pseudo-random series counterbalanced across the participants. For each participant, 99 final eccentricities were collected and the median was computed.

##### Apparatus

The apparatus was the same as in Experiment 1.

### Experiment 4 – Auditory Output

Here, participants were submitted to auditory stimuli generated with the dynamic characteristics of the motor performances recorded in experiment 2 and were asked to modify this dynamic to evoke a friction sound that conveys the most representative elliptic movement exclusively from the auditory modality. By using monophonic sounds, only dynamic information was transmitted to the participants, as opposed to the visual cases in which both spatial and dynamic cues were contained in the stimuli. Hence, this experiment intended to show whether geometric prototypes could emerge from the auditory stimulations, which would signify that spatial information is not strictly needed to evoke a prototype.

#### Methods

##### Participants

Twenty participants (9 women) with an average age of 32.1 years (SD = 9.56) voluntary took part in the experiment. None of them participated to the experiments 1, 2 & 3.

##### Stimuli

Stimuli were monophonic synthetic friction sounds simulating the motion of a pencil on a rough surface, e.g., a piece of paper. A physically based model which simulates the physical sound source as the result of successive impacts of a plectrum, here a pencil nail, on the asperities of a surface, was used (Conan et al., 2014). The surface roughness is modeled by a noise reflecting the different heights of the surface asperities. Movements on this surface is then simulated processing this noise according to the velocity profile of the movement: the faster the movement the more impacts there are. From a signal processing point of view, such an operation is equivalent to a low-pass filter on the noise with a central frequency directly linked to the movement velocity. The filtered noise is finally convolved with an impulse response simulating the resonance of an object such as a table. To control the synthesis model, we used the 60 motions (3 recordings x 20 participants) recorded in the motor experiment (Experiment 2).

##### Task

The participants were asked to modify the sound to evoke an elliptic motion for which the trajectory was the most representative of an ellipse. They were implicitly adjusting the delay between the recorded oscillators which implicitly modify the ellipse’s roundness. For each of the 60 movements to adjust, 4 periods of the initial recording were used to generate the stimuli. The participants were informed when they reached limits, i.e. 0 or 1 eccentricity, and the adjustment protocol was inspired by one proposed by Carlyon and colleagues (Carlyon et al., 2010). The eccentricity of the ellipse could be modified either with a large or a small modification of the actual eccentricity by using top-down and left-right-arrows respectively, by steps of .10/.3 and .05/.2 respectively. In order to avoid an experimental bias, for each opposite key, e.g., left/right, the roundness was modified with a different step so that participants could not count the number of taps to adjust the friction sound. The participants were prompted to explore the whole range of possibilities with the large arrows and to refine their adjustment with the smaller ones.

##### Apparatus

Sounds were presented through Sennheiser HD650 headphones and the sample rate of the soundcard was 44100 Hz with 16-bit resolution. The friction sounds were real-time synthesized with MATLAB software. The experiment was carried out in a quiet room. The sound was modified by the participants using the keyboard of the computer.

##### Data analysis

For each participant, 60 eccentricities values were collected and their median was computed to characterize the prototypical ellipse.

## Results

### Coherence between prototypes

We first investigated whether the four different experiments led to four different prototypes. For each experiment, we ensured the normality of the different distributions by using a Lilliefors test (all non-significant, Bonferroni corrected α=.05/4=.0125). It must be noted that for the auditory experiment the Lilliefors test was only marginally significant because of Bonferroni corrections (p=.0176). Therefore, in the following we used both parametric and non-parametric approaches. Two statistical tests were applied: (1) a non-parametric Kruskal-Wallis test with experiment as a factor (df=3; χ^2^_=_ 1.14; p=.76); (2) parametric Bonferroni corrected pairwise t-tests between the different experiments. Results were all non-significant (*df*=19, motor vs. visual static: p=.33, BF_10_=.35; motor vs. visual dynamic: p=.84, BF_10_=.23; motor vs. auditory: p=.92, BF_10_=.23; visual static vs. visual dynamic: p=.55, BF_10_=.27; visual static vs. auditory: p=.25, BF_10_=.42; visual dynamic vs. auditory: p=.82, BF_10_=.24) (Figure 2a). These tests thus supported an absence of significant difference between the motor, the visual - static and dynamic - and the auditory prototypes. *Eccentricity of the prototypes*. Secondly, we investigated the eccentricity of the prototypes for each experiment. The mean eccentricity values were: 1) motor: Mdn=.89, iqr=.067, M=.89, SD=.046; 2) visual static: Mdn=.91, iqr=.062, M=.90, SD=.044; 3) visual dynamic: Mdn=.91, iqr=.121, M=.88, SD=.086; 4) auditory: Mdn=.88, iqr=.049, M=.88, SD=.061 (Figure 2a). We determined a statistically based confidence interval for each prototype, one in each experiment. We linearly sampled the eccentricity range between 0 and 1 with 1000 equally spaced eccentricity values. We then ran 1000 one sample t-tests between the data distribution of each experiment and the 1000 eccentricity values. The results of each of the 1000 tests characterizes whether the distribution is different from the candidate eccentricity. For each of the 1000 t-tests, the inverse of the bayes factor BF_10,inverse_=1/BF_10_ was stored. BF_10,inverse_ can be interpreted exactly as BF_10_, but for *no difference* with the null hypothesis, the higher the BF_10,inverse_ the less different the distribution with the candidate eccentricity. We here assume that BF_10,inverse_>3 characterizes a fair threshold for no difference with the null hypothesis, i.e. the eccentricity value can be considered in the prototype range (Jeffreys, 1961; Goodman, 1999; Held & Ott, 2016). As results distributions were globally localized, the 1/BF_10_ curve is an inverse U-shape. For each experiment, we therefore used BF_10,inverse_=3 as a threshold to determine the lower and upper bound of the confidence interval. We also computed the eccentricity ecc_max_ leading to the highest BF_10,inverse_. This provides a third estimation of the elliptic prototype eccentricity for each experiment. This analysis provides the following results: motor: ecc_max_=.883, CI_BF10,inverse_=[.869,.897] - visual static: ecc_max_=.899, CI_BF10,inverse_=[.892,.909] - visual dynamic: ecc_max_=.888, CI_BF10,inverse_=[.871,.905] - auditory: ecc_max_=.880, CI_BF10,inverse_=[.870,.893]. To investigate specifically whether they differ from the known preferential ellipse of eccentricity .91, t-tests between the distributions and the .91 value were conducted. Results revealed no significant differences (*df*=19, Bonferroni corrections α=.05/4=.0125, motor: p=.1, BF_10_=.82 – visual static: p=.34, BF_10_=.354 – visual dynamic: p=.25, BF_10_=.42 - auditory: p=.05, BF_10_=1.37). Altogether, these analyses revealed that the experiments provide comparable prototypes, with an average eccentricity of .887. These prototypes are slightly below the preferential intermediate ellipse of eccentricity .91 observed in the literature (see General Discussion).

**Figure 2.**
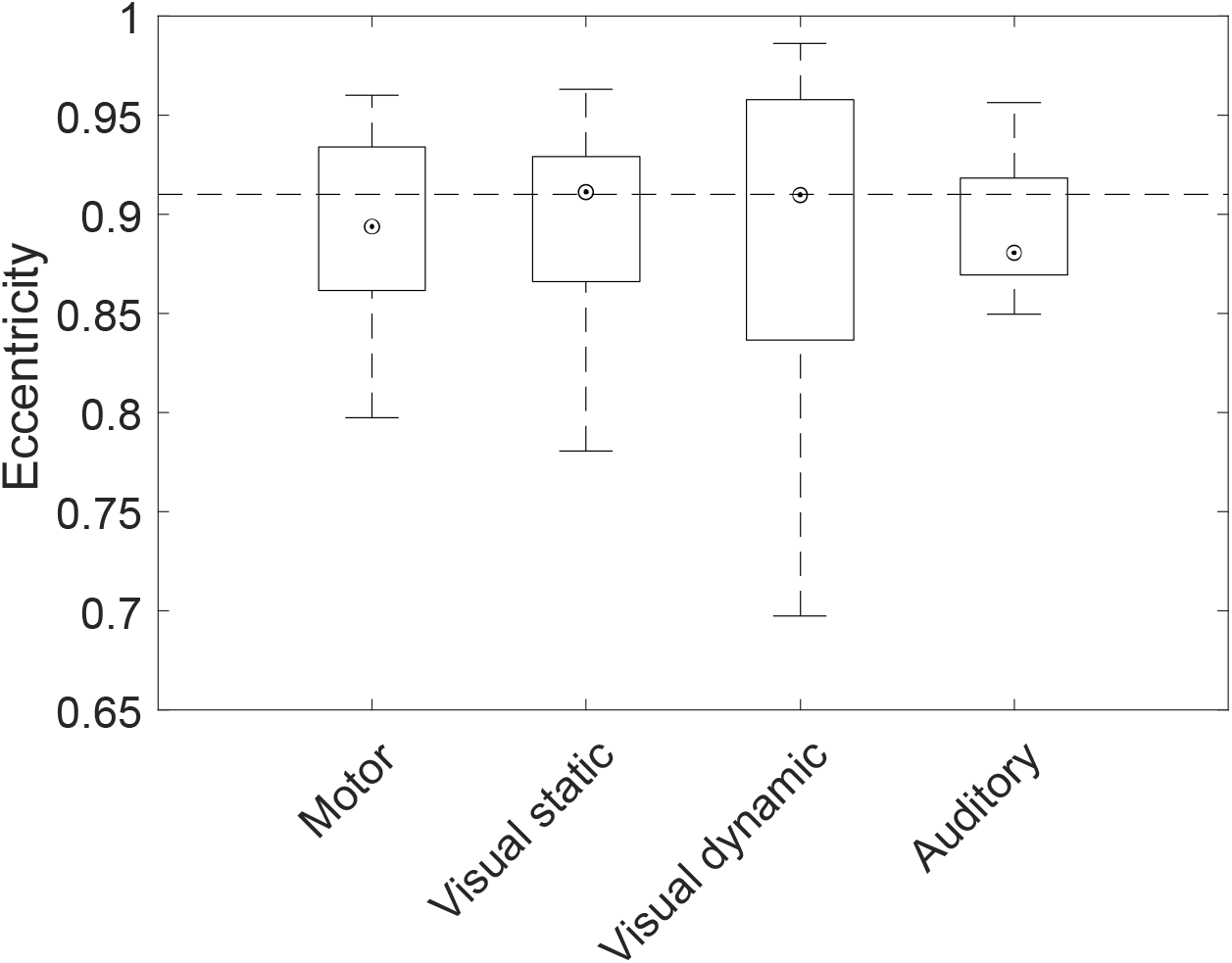
Averaged eccentricities obtained from the four experiments. The boxplot represents the median (dot), the 25^th^ and 75^th^ percentiles (the bottom and top edges of the box). The whiskers (vertical dashed line) extend to the most extreme data points below 1.5 the interquartile range. The dashed line indicates the value .91 corresponding to the elliptic movement attractor.

## General Discussion

In this study, we aimed at investigating whether a common prototype of elliptic shapes elicited across modalities, especially from auditory stimuli, could be identified. We compared the restituted geometry through different outputs (visual, motor and auditory) across a series of four experiments. A common elliptic prototype characterized by an eccentricity of .887 was uncovered, hence demonstrating the common encoding of this prototype.

This complements knowledge from previous studies on the existence of prototypical geometric shapes (Rosch, 1973, 1975; Feldman, 2000; Kalénine et al., 2013). This is also in line with those from Wamain et al. (2011), who highlighted that the ellipse drawn spontaneously has a relative phase slightly below 45°, i.e. an eccentricity slightly below.91, and with those from pure motor experiments aiming at revealing the preferentially drawn ellipse (Dounskaia et al., 2000; Sallagoïty et al., 2004; Danna et al., 2011) corresponding to stable states, so-called motor attractors (Kelso, 1986) (cf. Figure 2). Our results showed that the perceptual prototype – either visual or auditory – is coherent with the motor one, with an eccentricity value slightly below the intermediate ellipse of .91. This eccentricity difference may arise from the intention of the participants which were different because of a different task. Here we asked them to draw the most prototypical ellipse, which wasn’t present at all in the spontaneously drawn ellipses asked in Wamain et al. (2011). This slight difference suggests that we cannot obviously exclude that other cue also might have shaped the geometry of the prototype than motor constraint such as visual experience. Interestingly, when superimposing the prototypical ellipse obtained in the present study with the prototypical rectangle highlighted by Kalénine et al. (2013), common proportions can be observed. The length to width ratio for prototypical rectangles and the semi-major to semi-minor axis ratio for prototypical ellipses are both close to 2.3, which corresponds to .9 in term of eccentricity.

Above all, these results suggest a common coding of prototypical shape geometry and more generally further support the role of the sensorimotor loop on our perceptual processes (O’Regan and Noë, 2001; Varela et al., 1992; O’Regan, 2011). Interestingly, neuropsychological studies demonstrated that motor schemes are re-activated during perceptual processes through different sensory modalities (Chao and Martin, 2000; Creem-Regehr and Lee, 2005; Grafton et al., 1997; Grèzes and Decety, 2002; Kohler et al., 2002; Bangert et al., 2006; Zatorre et al., 2007), and through motor imagery (for a review see Jeannerod, 1995 or Grosprêtre, Ruffino and Lebon, 2015). Concerning handwriting, it has been shown that seeing a letter activates the cortical processes involved when producing the corresponding script (Longcamp et al., 2003; James and Gauthier, 2006; Longcamp et al., 2006). Similarly, perceiving a geometric shape may share processes involved when we are drawing it. In Experiment 4, an elliptic prototype that did not differ from the visual and motor outputs was elicited from monophonic sounds that only carried dynamic cues. This salient result shows that the elliptic prototype is not only geometric but also can be transmitted through a dynamic dimension. Hence without rejecting the necessity of visual experience in the emergence of prototypical shapes (Theurel et al., 2011; Kalénine et al., 2011), our data suggest that the prototypical shape is encoded through a common process based on the underlying covariation between biological kinematics and shape geometry which characterizes the corresponding drawing movement (Lacquaniti et al., 1983) and its perception through several sensory channels (Viviani et al., 1992; Viviani and Stucchi, 1989; Viviani et al., 1987; Thoret et al., 2014). This unified percept (Hommel et al., 2001) can then be recalled even through the auditory modality, which does not provide any spatial cues. Further experiments using the same auditory based paradigm, in particular with other geometric shapes, are needed to precisely assess the central role of the sensorimotor loop in the emergence of prototypical shape geometries.

## Limitations & Perspectives

This study has mainly two methodological limitations. Firstly the investigation was made at group level and subjects were different in each experiment. This allowed us to strikingly highlight the existence of a common prototype between a set of different participants in each modality. This was the main hypothesis of this study and the reason why the elliptic prototype was primarily investigated at group level. It would be of interest to investigate this issue at subject level to exclude that such a prototype may vary across modalities for a given set of participants. However, such a subject level investigation would require a different experimental procedure as one experiment might influence the result of the others. In this case it would not be possible to completely disentangle a potential priming between them. Secondly, the experiment was performed in controlled laboratory experimental conditions with a population of N=80 (4×20). The generalizability of the result could be improved by running such a series of experiments on a larger number of participants with online testing platforms. Lastly, and this is probably the most important conceptual limitation of this study, we have here investigated the prototype of only one shape, the ellipse, which is a very specific shape without any discontinuities, and which moreover can be drawn continuously. To generalize our results, it would be relevant to run the experiment with a more complete set of shapes such as rectangles or triangles, that have been subject to a certain number of studies in the past. This would nevertheless require refinements in the sound modeling procedure due to the discontinuities in the drawing movement of such shapes.

## Supporting information

Matlab script to regenarate figure and results

## Authors contributions

Conceived and designed the experiments: ET MA LB SY RKM. Performed the experiments: ET. Analyzed the data: ET MA LB SY RKM. Contributed reagents/materials/analysis tools: ET. Wrote the paper: ET MA LB SY RKM

## Acknowledgments

This work was funded by the French National Research Agency (ANR) under the MetaSon: Métaphores Sonores (Sound Metaphors) project (ANR-10-CORD-0003) in the CONTINT 2010 framework and the SoniMove Project (ANR-14-CE24-0018). E.T. was funded through an ILCB/BLRI grant no. ANR-16-CONV-0002 (ILCB), ANR-11-LABX-0036 (BLRI) and the Excellence Initiative of Aix-Marseille University (A*MIDEX). The authors would like to thank Thomas Bordonné and Joris Agator for their help running the experiments. E.T. is thankful to Jeremy Danna for his thoughtful suggestions and guidance at the very beginning of this project.

