## Supplementary material for "Hearing elliptic movements reveals the imprint of action on prototypical geometries": Matlab script to regenarate figure and results: analysis_cognition.m

```
%
% generate results and figure 2 of the paper
% Thoret, E. et al. (2023) Hearing elliptic movements reveals the imprint of action on prototypical geometries. Cognition
%
% The following script needs the following package for bayes factor analysis
% https://github.com/klabhub/bayesFactor
%
% results are logged in 'log_results.txt'
%

clearvars;

%% load data from the auditory experiment
path_ = './' ;
addpath(genpath(path_)) ;
filelist = dir([path_ 'aud_*.mat']) ;
listSubject = 1:length(filelist) ;
nbSubjects = length(listSubject) ;

resultsAllSubjects = zeros(nbSubjects,60) ;
resultMeanSubj = [] ;

for iSubject = listSubject
    load([path_ filelist(iSubject).name]);
    disp(filelist(iSubject).name) ;
    resultOneSubject = zeros(1,60) ;
    for i_trial = 1:60
        temp = phi0_struct{i_trial} ;
        resultOneSubject(i_trial) = temp(end) ;
    end
    resultOneSubject_e = sqrt(abs(1.-tan(resultOneSubject/2).^2)) ;
    resultsAllSubjects(iSubject,:) = resultOneSubject_e ;
end

eccentricities_all = resultsAllSubjects ;
relativePhase_all  = 2*atan(sqrt(1-resultsAllSubjects.^2))*180/pi ;

%% load data from the motor, visual static, and visual dynamic experiments
ecc_auditory = eccentricities_all ;
load('tabT.mat','tabT','ecc_tab') ;
ecc_motor = ecc_tab ;
load('results_visual_static.mat','ecc_f') ;
ecc_visual_static = sqrt(1-tan(pi*ecc_f/2/180).^2) ;
load('results_visual_dynamic.mat','ecc_f') ;
ecc_visual_dynamic = sqrt(1-tan(pi*ecc_f/2/180).^2) ;
clearvars ecc_tab ;

% subject medians of data aggregated in one array 20 x 4
data = [median(reshape(ecc_motor,20,3),2) (median(ecc_visual_static,2)) (median(ecc_visual_dynamic,2)) (median(ecc_auditory,2))] ;

median_results = [median(data) ; iqr(data)] ; % medians & iqr
mean_results = [mean(data) ; std(data)] ; % means & std

%% normality tests
[h1, p1] = lillietest(data(:,1)) ;
[h2, p2] = lillietest(data(:,2)) ;
[h3, p3] = lillietest(data(:,3)) ;
[h4, p4] = lillietest(data(:,4)) ;

lilliefors_p_val = [p1, p2, p3, p4] ; % lilliefors stats

%% figure 2
ecc_motor_tabStat = median(reshape(ecc_motor,3,20),1)' ;
ecc_vs_tabStat    = median(ecc_visual_static,2) ;
ecc_vd_tabStat    = median(ecc_visual_dynamic,2) ;
ecc_aud_tabstat   = median(ecc_auditory,2) ;

tab = [ecc_motor_tabStat ecc_vs_tabStat, ecc_vd_tabStat, ecc_aud_tabstat] ;

boxplot(tab,'BoxStyle','outline','colors','k','MedianStyle','target','outliersize',0.001,...
                'Widths',.5, 'symbol', '')
axis([0 5 .65 1])
XTick_pos = (1:4);
XTick_labels = {'Motor', 'Visual static', 'Visual dynamic', 'Auditory'} ;
xtickangle(45)
hold on
plot(linspace(0,5,24),ones(1,24)*0.91,'--k','linewidth',.5)
set(gca, 'XTick', XTick_pos, 'XTickLabel', XTick_labels,'fontsize',18); % 10 ticks 
ylabel('Eccentricity')
hold off

%% stats
% kruskal-wallis
[p, tbl, stats] = kruskalwallis(tab) ;
close all;
% t-test vs .91
ecc = .91 ;
[bf10_motor_91,p_motor]       = bf.ttest(tab(:,1),ecc) ;
[bf10_visstat_91,p_visstat]   = bf.ttest(tab(:,2),ecc) ;
[bf10_visdyn_91,p_visdyn]     = bf.ttest(tab(:,3),ecc) ;
[bf10_auditory_91,p_auditory] = bf.ttest(tab(:,4),ecc) ;

%% BF10 confidence interval for each modality
tab_ress = zeros(4,3) ;

[maxBF10, ci_low, ci_up, bf10_tab_1, ~] = ...
    bf10_CI(tab(:,1), [.0 1], 3, 1) ;
tab_ress(1,1) = maxBF10 ;
tab_ress(1,2) = ci_low  ;
tab_ress(1,3) = ci_up   ; 

[maxBF10, ci_low, ci_up, bf10_tab_2, ~] = ...
    bf10_CI(tab(:,2), [.0 1], 3, 1) ;
tab_ress(2,1) = maxBF10 ;
tab_ress(2,2) = ci_low  ;
tab_ress(2,3) = ci_up   ; 

[maxBF10, ci_low, ci_up, bf10_tab_3, ~] = ...
    bf10_CI(tab(:,3), [.0 1], 3, 1) ;
tab_ress(3,1) = maxBF10 ;
tab_ress(3,2) = ci_low  ;
tab_ress(3,3) = ci_up   ; 

[maxBF10, ci_low, ci_up, bf10_tab_4, ~] = ...
    bf10_CI(tab(:,4), [.0 1], 3, 1) ;
tab_ress(4,1) = maxBF10 ;
tab_ress(4,2) = ci_low  ;
tab_ress(4,3) = ci_up   ; 

%% BF10 t-test for comparison between modalities
[bf10_m_vs,p_m_vs]   = bf.ttest(tab(:,1),tab(:,2)) ;
[bf10_m_vd,p_m_vd]   = bf.ttest(tab(:,1),tab(:,3)) ;
[bf10_m_a,p_m_a]    = bf.ttest(tab(:,1),tab(:,4)) ; 
[bf10_vs_vd,p_vs_vd]  = bf.ttest(tab(:,2),tab(:,3)) ; 
[bf10_vs_a,p_vs_a]   = bf.ttest(tab(:,2),tab(:,4)) ; 
[bf10_vd_a,p_vd_a]   = bf.ttest(tab(:,3),tab(:,4)) ; 

%% output results
clc
disp('Liliefors p-values:\n')
disp(lilliefors_p_val)
disp('Kruskal-Wallis\n')
disp(['    chi2=', num2str(cell2mat(tbl(2,5))), '; p=',num2str(cell2mat(tbl(2,6)))])
disp(' ')
disp('Paiwise t-test:')
disp(['    motor / vs: bf10=',num2str(bf10_m_vs),', p=',num2str(p_m_vs)])
disp(['    motor / vd: bf10=',num2str(bf10_m_vd),', p=',num2str(p_m_vs)])
disp(['    motor / auditory: bf10=',num2str(bf10_m_a),', p=',num2str(p_m_vs)])
disp(['    vs / vs: bf10=',num2str(bf10_vs_vd),', p=',num2str(p_m_vs)])
disp(['    vs / auditory: bf10=',num2str(bf10_vs_a),', p=',num2str(p_m_vs)])
disp(['    vd / vs: bf10=',num2str(bf10_vs_vd),', p=',num2str(p_m_vs)])
disp(' ')
disp('Median eccentricity:')
disp(median_results)
disp('Mean eccentricity:')
disp(mean_results)
disp('Eccentricy confidence interval obtained with BF10 analysis')
disp(['    motor: eccmax=', num2str(tab_ress(1,1)), ' , CI_BF10=[', num2str(tab_ress(1,2)), ',', num2str(tab_ress(1,3)),']']) ;
disp(['    visual static: eccmax=', num2str(tab_ress(2,1)), ' , CI_BF10=[', num2str(tab_ress(2,2)), ',', num2str(tab_ress(2,3)),']']) ;
disp(['    visual dynamic: eccmax=', num2str(tab_ress(3,1)), ' , CI_BF10=[', num2str(tab_ress(3,2)), ',', num2str(tab_ress(3,3)),']']) ;
disp(['    auditory: eccmax=', num2str(tab_ress(4,1)), ' , CI_BF10=[', num2str(tab_ress(4,2)), ',', num2str(tab_ress(4,3)),']']) ;
disp(' ')
disp('t-test vs. .91:')
disp(['    motor: bf10=',num2str(bf10_motor_91),', p=',num2str(p_motor)])
disp(['    visual static: bf10=',num2str(bf10_visstat_91),', p=',num2str(p_visstat)])
disp(['    visual dynamic: bf10=',num2str(bf10_visdyn_91),', p=',num2str(p_visdyn)])
disp(['    auditory: bf10=',num2str(bf10_auditory_91),', p=',num2str(p_auditory)])
disp(' ')
disp('Average eccentricity')
disp(['    M=',num2str(mean(tab_ress(:,1)))])

%% write results
fid=fopen('./log_results.txt','w+');
fprintf(fid,'Motor | Visual static | Visual Dynamic | Auditory \n') ;
fprintf(fid,'\n');
fprintf(fid,'Liliefors p-values:\n') ;
fprintf(fid,['    ', num2str(lilliefors_p_val), '\n']) ;
fprintf(fid,'\n');
fprintf(fid,'Kruskal-Wallis');
fprintf(fid,['\n', '    chi2=', num2str(cell2mat(tbl(2,5))), '; p=',num2str(cell2mat(tbl(2,6))), '\n']);
fprintf(fid,'\n');
fprintf(fid,'Paiwise t-test:\n');
fprintf(fid,['    motor / vs: bf10=',num2str(bf10_m_vs),', p=',num2str(p_m_vs), '\n']);
fprintf(fid,['    motor / vd: bf10=',num2str(bf10_m_vd),', p=',num2str(p_m_vd), '\n']);
fprintf(fid,['    motor / auditory: bf10=',num2str(bf10_m_a),', p=',num2str(p_m_a), '\n']);
fprintf(fid,['    vs / vs: bf10=',num2str(bf10_vs_vd),', p=',num2str(p_vs_vd), '\n']);
fprintf(fid,['    vs / auditory: bf10=',num2str(bf10_vs_a),', p=',num2str(p_vs_a), '\n']);
fprintf(fid,['    vd / vs: bf10=',num2str(bf10_vs_vd),', p=',num2str(p_vs_vd), '\n']);
fprintf(fid,'\n');
fprintf(fid,'Median eccentricity:');
fprintf(fid,'\n');
fprintf(fid,['    ', num2str(median_results(1,:))]);
fprintf(fid,'\n');
fprintf(fid,['    ', num2str(median_results(2,:))]);
fprintf(fid,'\n');
fprintf(fid,'\n');
fprintf(fid,'Mean eccentricity:');
fprintf(fid,'\n');
fprintf(fid,['    ', num2str(mean_results(1,:))]);
fprintf(fid,'\n');
fprintf(fid,['    ', num2str(mean_results(2,:))]);
fprintf(fid,'\n');
fprintf(fid,'\n');
fprintf(fid,'Eccentricy confidence interval obtained with BF10 analysis\n');
fprintf(fid,['    motor: eccmax=', num2str(tab_ress(1,1)), ' , CI_BF10=[', num2str(tab_ress(1,2)), ',', num2str(tab_ress(1,3)),']', '\n']) ;
fprintf(fid,['    visual static: eccmax=', num2str(tab_ress(2,1)), ' , CI_BF10=[', num2str(tab_ress(2,2)), ',', num2str(tab_ress(2,3)),']', '\n']) ;
fprintf(fid,['    visual dynamic: eccmax=', num2str(tab_ress(3,1)), ' , CI_BF10=[', num2str(tab_ress(3,2)), ',', num2str(tab_ress(3,3)),']', '\n']) ;
fprintf(fid,['    auditory: eccmax=', num2str(tab_ress(4,1)), ' , CI_BF10=[', num2str(tab_ress(4,2)), ',', num2str(tab_ress(4,3)),']', '\n']) ;
fprintf(fid,'\n');
fprintf(fid,'t-test vs. .91:\n');
fprintf(fid,['    motor: bf10=',num2str(bf10_motor_91),', p=',num2str(p_motor), '\n']);
fprintf(fid,['    visual static: bf10=',num2str(bf10_visstat_91),', p=',num2str(p_visstat), '\n']);
fprintf(fid,['    visual dynamic: bf10=',num2str(bf10_visdyn_91),', p=',num2str(p_visdyn), '\n']);
fprintf(fid,['    auditory: bf10=',num2str(bf10_auditory_91),', p=',num2str(p_auditory), '\n']);
fprintf(fid,'\n');
fprintf(fid,'Average eccentricity:\n');
fprintf(fid,['    M=',num2str(mean(tab_ress(:,1)))]);
fclose(fid); 

%% function to determine the confidence interval based on BF10
function [maxBF10, ci_low, ci_up, bf10_tab, p_tab] = bf10_CI(data, range, threshold_BF10, inverse)
    bf10_tab = [] ;
    p_tab = [] ;
    lin_ecc = linspace(range(1),range(2),1000) ;
    for ecc=lin_ecc
        [bf10,p] = bf.ttest(data,ecc) ;
        bf10_tab = [bf10_tab; bf10];
        p_tab = [p_tab; p] ;
    end
    if inverse == 1
        bf10_tab = 1./bf10_tab ;
    end
    indMin = min(find((bf10_tab)>threshold_BF10)) ;
    indMax = max(find((bf10_tab)>threshold_BF10)) ;
    [~,indMaxValue] = max(bf10_tab) ;

    ci_low  = lin_ecc(indMin) ;
    ci_up   = lin_ecc(indMax) ;
    maxBF10 = lin_ecc(indMaxValue) ;

end
```
